# Molecular and cellular signatures differentiate Parkinson’s disease from Parkinson’s disease with dementia

**DOI:** 10.1101/2025.03.04.641379

**Authors:** Aine Fairbrother-Browne, Melissa Grant-Peters, Jonathan W. Brenton, Hemanth Nelvagal, Regina Reynolds, Yau Mun Lim, Emil K Gustavsson, Hannah Macpherson, Kylie Montgomery, James R. Evans, Amy R Hicks, Nancy Chiraki, Toby Curless, Raquel Real, Theodoros Xenakis, Henry Houlden, Huw R Morris, Sonia Gandhi, Nicholas W. Wood, John Hardy, Zane Jaunmuktane, Mina Ryten

## Abstract

Parkinson’s disease (PD) affects millions of people worldwide, and up to 40% of these patients develop dementia, profoundly affecting their quality of life. Whether Parkinson’s disease dementia (PDD) simply represents a late stage of PD or constitutes a distinct neurodegenerative process remains unresolved. To clarify this, we generated the largest single nuclear transcriptomic atlas of PD and PDD to date—almost one million nuclei derived from the anterior cingulate cortex and inferior parietal lobule of 64 post-mortem donors. By integrating these data with long-read RNA-seq, we found that the cellular compositions, biological pathways, and molecular profiles diverge substantially between PD and PDD, with limited overlap in differentially expressed genes and pathways. While PD was characterised by widespread upregulation of gene expression programs and robust regional signatures, PDD showed extensive pathway downregulation, loss of cortical regional identity, and significant shifts in transcript usage, including alterations in *APP* isoforms that may influence pathological amyloid beta accumulation. These findings reveal that PD and PDD represent fundamentally distinct disease states, offering important insights for understanding their underlying mechanisms and will guide the development of targeted therapies and more effective clinical trials.

## Introduction

Parkinson’s disease (PD) is a progressive neurodegenerative disorder characterized by an initial loss of dopaminergic neurons in the substantia nigra pars compacta, followed by other brain regions, and the presence of Lewy pathology. While motor symptoms are primary and respond to dopamine replacement therapy, there has been increasing recognition of the importance of non-motor symptoms which are generally difficult to treat. Of the non-motor symptoms, dementia is the most important, and is a major determinant of quality of life.

Between 20-40% of patients with PD develop dementia, with males at higher risk^1,2^. Furthermore, the risk of dementia increases over time: in longitudinal clinical studies, 26% of patients developed dementia within three years of a PD diagnosis^3^, but after 20 years 83% of patients had developed dementia ^4^. Diagnosis depends on the timing of symptom onset, namely if dementia precedes or appears within a year of motor symptoms, it is classified as dementia with Lewy bodies (DLB), otherwise it is termed Parkinson’s disease dementia (PDD)^5,6^.

It remains unclear whether PDD is simply a more advanced form of PD, driven by shared molecular processes, or a distinct disease entity with differing biology. Despite notable clinical differences between PD and PDD, they cannot yet be confidently distinguished pathologically, as both have Lewy pathology and may also exhibit co-pathologies, including hyperphosphorylated tau (pTau) and amyloid beta (Aβ), hallmarks of Alzheimer’s disease (AD)^7–9^. Therefore, the relationships between the development of dementia in PD and the underlying molecular features of disease, including neuropathology, remain unclear.

Recently, there have been efforts to address this through the identification of genetic risk factors associated specifically with dementia risk in PD. These studies suggest that the genetic architecture of progression to dementia differs from PD causation. For example, while risk variants of *GBA1* are known to increase the risk of both PD and PDD, rare variants in *LRRK2* are strongly associated with PD and may protect against progression to dementia^10–12^. In addition, genome-wide association studies have highlighted the association of PDD and DLB with AD-related genes, such as *BIN1*, *LRP1B* and *APOE,* but not with PD-related genes^13,14^.

Understanding the molecular differences between PDD and PD is essential for improving prognosis, patient stratification, and therapeutic development. However, the lack of reliable biomarkers to predict dementia risk in PD hampers these efforts, often grouping biologically distinct patients in clinical trials and confounding drug responses. As exemplified by amyotrophic lateral sclerosis (ALS) and central nervous system (CNS) demyelinating disorders, drug development has benefited from patient stratification based on a molecular understanding of disease state, such as the underlying mutations causing disease (ALS) (Miller et al. 2022) or antibodies specific to a clinical group (CNS demyelination)^15–17^. To address this knowledge gap for PD, we used single-nucleus RNA sequencing (snRNAseq) and long-read RNA sequencing to profile human brain tissue derived from patients with PD and PDD. Our analysis revealed distinct molecular and cellular features, particularly in excitatory neurons.

## Results

### Cortical layer 4 RORB neurons are less represented in PD than in controls

Our study cohort comprises a snRNAseq data set of almost 1 million nuclei from 64 participants (950,156 nuclei). These were generated from three clinical groups: Parkinson’s disease (PD, n = 21), Parkinson’s disease with dementia (PDD, n = 21) and controls without neurological disease (Control, n = 22) (**Fig 1A**). For all participants, both clinical notes and neuropathology were reviewed to ensure diagnostic accuracy. The mean age at death across groups ranged between 76 years (PDD) and 87 years old (Controls), with a slightly higher proportion of male samples among PDD (female/male ratio: 1:2.5) and PD (1:1.33) affected individuals (**Fig 1B**), reflecting the higher prevalence of PD and PDD in men when compared to women. Median disease length was also comparable for both disease groups (MPD_length = 11; MPDD_length = 10 years). For each individual, samples were taken from two cortical brain regions, namely the anterior cingulate gyrus (ACG) and the inferior parietal lobule (IPL). All PD and PDD participants were diagnosed with late stage Parkinson’s disease (Braak stages 5 and 6) and the two cortical regions were chosen for their differential neuropathology, with ACG having a higher Lewy body load than IPL at Braak stages 5 and 6. This experimental design provided a pseudo-temporal analytical framework to study disease progress between individuals and within individuals. We also determined the α-synuclein and Aβ pathological load (**Fig 1C,D**).

**Figure 1.**
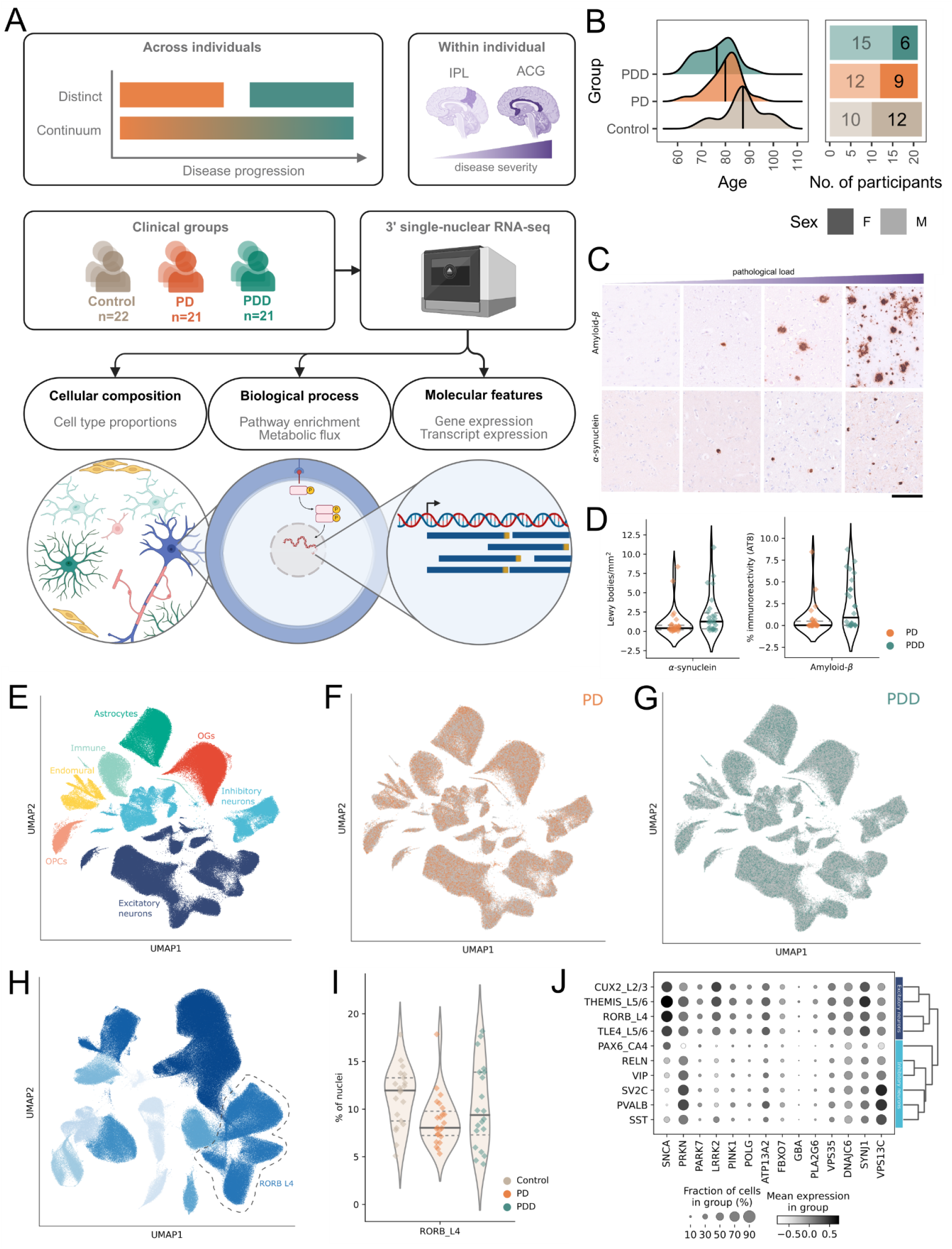
In-depth molecular and cellular comparison of PD and PDD. A. To test whether Parkinson’s disease without cognitive impairment (PD) and Parkinson’s disease with dementia (PDD) are distinct disease processes or whether they are part of the same disease continuum, we performed single nucleus RNA sequencing on post-mortem brain samples from the ACG) and inferior parietal lobule (IPL) from 64 individuals (controls n = 22; PD n = 21; PDD n = 21). This strategy provided us with an approximation of disease progression for each condition, since the ACG has more severe pathology than the IPL. We leveraged this data to profile the cellular composition, biological processes and molecular features in controls, PD and PDD. B. The mean age at death across groups ranged between 76 years (PDD) and 87 years old (controls), with a slightly higher proportion of male samples among PDD (female/male ratio: 1:2.5) and PD (1:1.33) affected individuals. C. The spectrum of α-synuclein and Aβ pathology density in the cortex of the superior frontal gyrus across controls, PD and PDD. From left to right, images show increasing density of pathological misfolded α-synuclein and Aβ aggregates. The top row corresponds to Aβ pathology composed of diffuse deposits and places with central cores. The bottom row corresponds to α-synuclein immunoreactive Lewy pathology composed of Lewy bodies and Lewy neurites. The scale bar shown in the bottom right corresponds to 100µm. D. PD and PDD quantitative pathology of Aβ and α-synuclein load in the superior frontal gyrus across controls. E. Our single-nucleus RNA-sequencing dataset included 950,156 nuclei from individuals with PD, PDD and controls with no neurological conditions. Following QC and cell type annotation, we had 789,708 nuclei represented across 7 major cell types, as determined by data-driven clustering, leiden community detection and manual annotation. F. Our final data set contained 267,150 nuclei from PD, which were represented across all annotated clusters. G. Our final data set contained 243,602 nuclei from PDD, which were represented across all annotated clusters. H. Clustering and annotation of neuronal cell types subdivided this population into 10 neuronal subtypes, of which four are excitatory and map to specific layers of the cortex. There were six further neuronal populations of inhibitory neurons. We highlight here the RORB_L4 population which predominantly resides in cortical layer 4. I. Cell type proportion analysis shows that RORB_L4 excitatory neuron difference between controls and PD is significant (FDR-adjusted p-value = 7.39×10^-3^). This cell type also had a downward trend in PDD, but this is not statistically significant (FDR-adjusted p-value = 8.29×10^-1^). J. Here we show the variation of PD-associated normalised gene expression across neuronal populations. The majority of genes are expressed more highly in excitatory neurons, such as *SNCA*, *LRRK2*, *PINK1*, *SYNJ1*. Others, such as *PRKN* and *VPS13C* are more highly expressed in inhibitory neurons. Notably, hierarchical clustering of the expression of these genes distinguishes excitatory and neuronal populations.

All samples underwent nuclear isolation and snRNAseq. Following removal of doublets and low quality cells, a total of 789,708 nuclei remained, capturing on average 5,850 nuclei per sample, thus enabling robust cell-type specific analyses (**Supp Fig 1A,B**). Similar numbers of nuclei were captured across the ACG (n = 399,932) and IPL (n = 389,776), and across experimental groups (controls, n = 278,956; PD, n = 267,150; and PDD, n = 243,602; **Supp Fig 1C**).

Following batch correction, dimensionality reduction and Leiden community detection, the nuclei were divided into seven major cell compartments including excitatory neurons, inhibitory neurons, oligodendrocytes, oligodendrocyte precursor cells (OPCs), astrocytes, and endomural - which contains both endothelial and mural cells - and immune cells (**Fig 1E**). Given the large number of nuclei captured from each sample, major cell classes were further partitioned into cell types and states, resulting in a total of 22 clusters. All clusters were distributed across each experimental group and tissue, meaning that none were disease-specific (**Fig 1F,G**), as might be expected for a late-onset disorder without a neurodevelopmental component. Clusters were manually annotated, identifying two oligodendrocyte (mature oligodendrocytes and oligodendrocyte progenitor cells), three astrocyte (CD44-high, RBFOX1-positive, SLC1A2- and MERTK-high), three immune (microglia, T-cells and B-cells), five endomural (fibroblasts, pericytes, endothelial cells, venous endothelia and smooth muscle cells), five inhibitory neuron (SST, RELN, PVALB, SV2C, PAX6 CA4) and four excitatory neuron clusters. The annotation of the latter included their cortical layer of origin (CUX2 layer 2, RORB layer 4, TLE4 layer 5/6, THEMIS layer 5/6) (**Supp Fig 2**). Neurons were the most abundant cell type and accounted for 55.3% of all nuclei in this dataset (**Fig 1H**).

**Figure 2.**
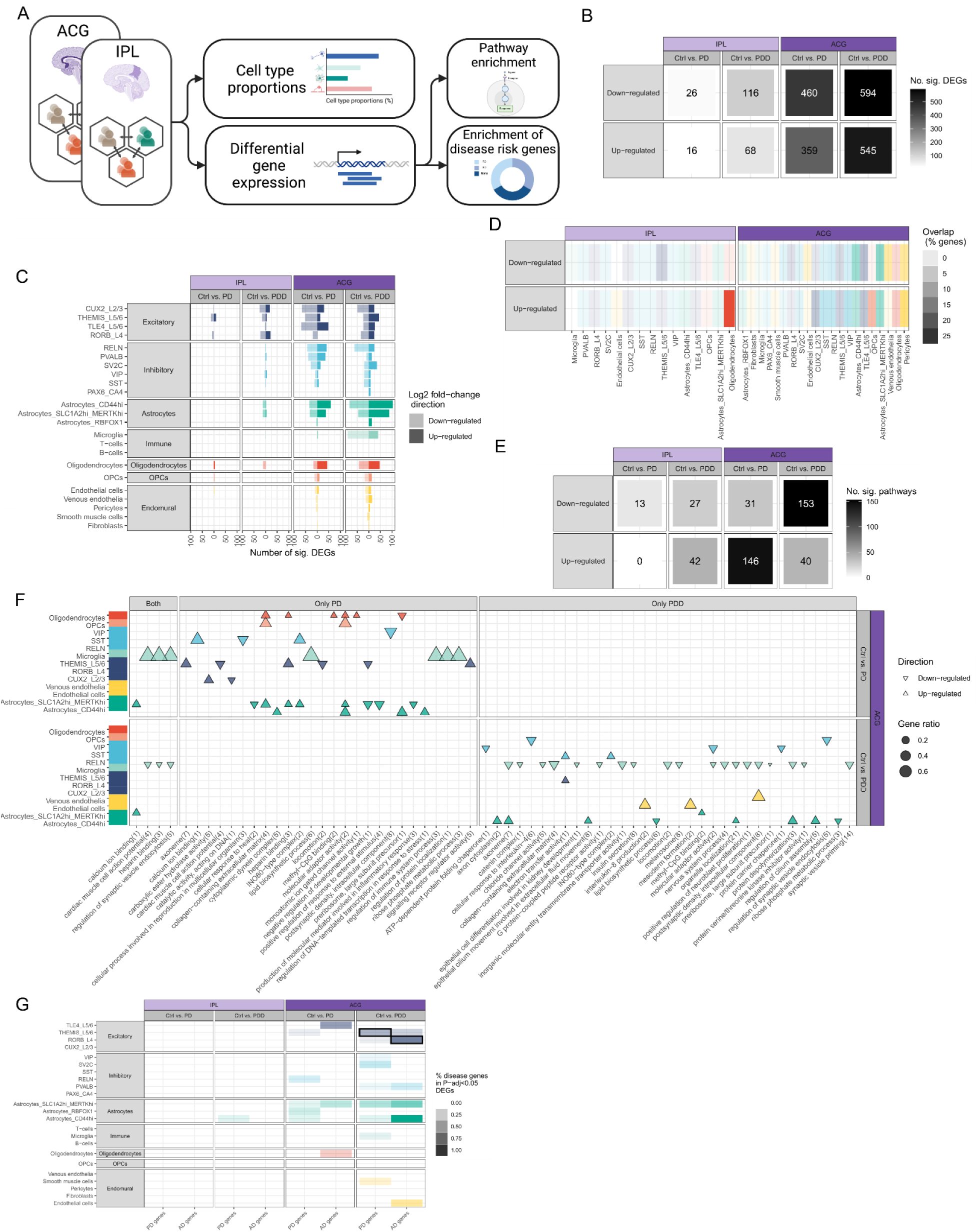
PD and PDD differential gene expression and pathway enrichment differ in nature and directionality. A. Workflow to summarise the structure of the inter-group comparison analyses. For each brain region we compared cell type proportions and differential gene expression in controls, PD and PDD. Using the list of differentially expressed genes, we assessed pathway enrichment and enrichment of genes associated with PD and Alzheimer’s disease risk. B. Heatmap to summarise the number of significant differentially expressed genes (DEGs) across tissues, group comparisons and DEG directionality (up- or down-regulated, FDR-corrected p-value <0.05). The counts represent the sum total of significant genes across all cell types. The ACG had most gene expression changes, as expected in light of higher pathological load of this region. PDD had more gene expression changes than PD for both brain regions. C. Bar plot showing total number of DEGs per cell type, stratified based on directionality. Gene expression changes were higher in the ACG than IPL for PD and PDD. PD had a higher number of DEGs in neuronal cell types, while PDD had more changes in glia - particularly microglia and astrocytes. D. Heatmap showing the number of overlapping PD and PDD DEGs. This is shown for both brain regions, stratified by directionality and cell type. E. Heatmap to summarise the number of significant pathways that are significantly enriched in DEGs for each brain region and group comparison (FDR-adjusted p-value < 0.05). The counts represent the total sum of significant pathways sum across all cell types. As expected based on DEGs, the majority of pathways are enriched in the ACG. PDD had predominantly down-regulated pathways (n = 153), while PD had predominantly up-regulated pathways (n = 146). F. Pathway enrichments in the ACG for PD and PDD across all cell types. The directionality of the pathway is encoded by triangle orientation (down-facing indicates down-regulation, upward facing indicates up-regulation) and the size of the triangle represents the gene ratio (proportion of genes in a pathway which are DEGs). Major cell types are indicated by colour. Most pathways were specific to PD or PDD, with only four pathways being enriched in both conditions and in the same cell type. PD pathway enrichment tended to be spread across multiple cell types, while PDD pathway enrichment was predominantly in microglia, inhibitory neurons and astrocytes. G. Heatmap showing hypergeometric enrichment of genes identified in GWAS studies for PD (n=183) and Alzheimer’s disease (n=111) among DEGs per brain region and cell type. Areas highlighted by red frame indicate significant enrichment, specifically, PDD DEGs in RORB_L4 neurons were enriched for AD-associated genes (FDR-adjusted p-value = 1.96 ×10^-2^) and THEMIS_L5/6 were enriched for genes associated with PD (FDR-adjusted p-value = 1.96 ×10^-2^).

Next, we compared cell type proportions across clinical groups relative to controls since disease-associated processes, such as cell death, could impact on tissue composition. We found that cell type proportion changes were most evident in PD (**Supp Fig 3**). Notably, in the ACG we found that RORB_L4 neurons, which account for 19.3% of all neurons, were less abundant in PD than in controls (FDR-adjusted p-value = 7.39×10^-3^) and this trend was also evident in PDD (**Fig 1I**). In common with this, some other excitatory neuronal cell types also expressed high levels of several genes causally-implicated in PD, including *SNCA*, *LRRK2* and *PINK1* (**Fig 1J**)^18^. Given that neurons cannot proliferate, this shift suggested the RORB_L4 neurons may be selectively vulnerable.

**Figure 3.**
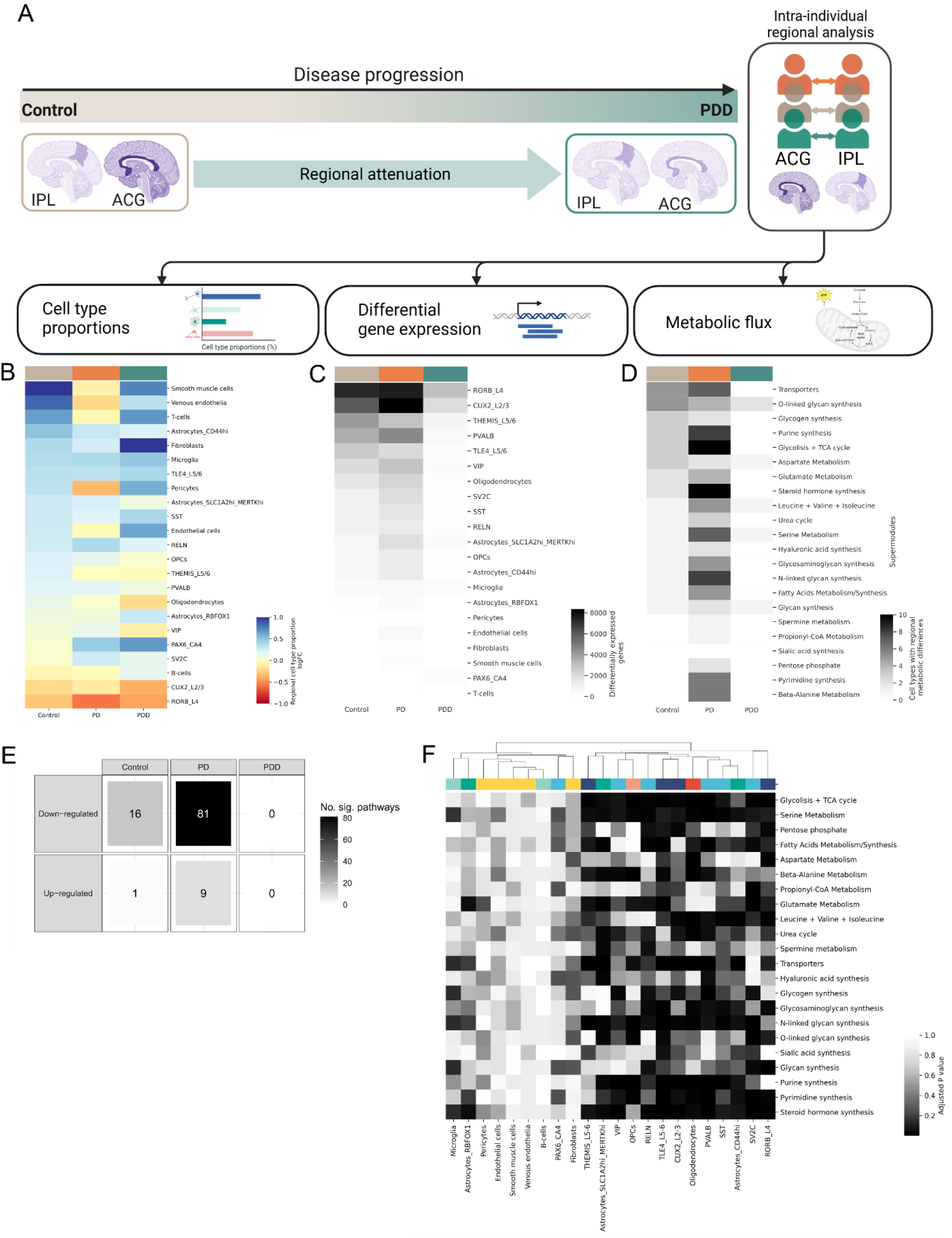
Regional attenuation is a feature of PDD, but not of PD. A. Schematic of regional attenuation hypothesis. Due to the large number of downregulated pathways in the PDD ACG, we hypothesised that this indicated a loss of region-specific features. To investigate this, we performed a cross-regional paired analysis. For this, we compared cell type proportion, differential gene expression and metabolic flux in the ACG in relation to IPL. B. We assessed the natural log-fold change of ACG cell type proportions relative to IPL in all groups. Cell type proportion changes in ACG relative to IPL show in a neurotypical state certain cell types tend to be more prevalent in the ACG (e.g. endomural, smooth muscle cells), while excitatory neurons (CUX2_L2/3, RORB_L4) are less prevalent in the ACG. In contrast with controls, PD had bidirectional changes across multiple cell types: RORB_L4 neurons were more depleted in ACG relative to IPL, smooth muscle cells and venous endothelia were less prevalent. In PDD, a depletion of RORB_L4 and CUX2_L2 neurons was present, albeit less pronounced than in PD. Most other cell types in PDD were proportionally more represented in the ACG than in the IPL, including fibroblasts, smooth muscle cells and venous endothelia. C. This heatmap shows the number of genes differentially expressed in the ACG when compared to the IPL. Here we show the total sum of DEGs across all cell types for each group. Differential gene expression of controls shows that RORB_L4 and CUX2_L2/3 neurons have the most regional specificity (i.e. the most differentially expressed genes). In PD, the number of differentially expressed genes between regions increases across all cell types, while in PDD there is a decrease in differentially expressed genes between regions. D. We inferred metabolic flux of 22 metabolic supermodules based on cell-specific gene expression and show in the heatmap a sum of the number of cell types with significant changes in the supermodule in ACG when compared to IPL (FDR-adjusted p-value <0.05). In controls, we found that certain metabolic flux supermodules varied between regions for a large number of cell types (e.g. transporters, O-linked glycan synthesis). In PD, the number of cell types with differing metabolic flux between regions increased substantially, with almost half of cell types with changes in the flux of glycolysis and TCA cycle. In PDD, regional metabolic flux differences were only present in 5 of the tested metabolic supermodules and only across 1-2 cell types, suggesting increased uniformity in metabolic flux across the tested brain regions E. Heatmap showing the total number of pathways differentially expressed between regions for each group. There were inherent differences in controls, mainly down-regulation. Regional differences were more pronounced in PD, with over 5 times more pathways down-regulated in PD than in controls. Meanwhile, PDD had no differential expression of pathways. F. Heatmap of the FDR-adjusted p-values testing metabolic differences across brain regions in PD. The tested cell types were hierarchically clustered to show which cell types had the most similar metabolic profiles. Clustering showed that there were two main clusters: one with major metabolic differences between regions, consisting of populations of neurons and oligodendrocytes, and one with fewer metabolic differences across between regions, consisting of immune cells, endomural cells and subsets of astrocyte populations.

### PD and PDD differ in nature and directionality of differentially expressed genes and pathways

Next, we performed differential gene expression analysis by cell type to further characterise the cell types and biological processes associated with each clinical group (**Fig 2A**). Gene counts were pseudobulked by sample and cell type using the statistically and computationally robust analytical framework provided by the tool dreamlet^19^. Consistent with the higher load of Lewy pathology in ACG as compared to IPL, we identified more differentially expressed genes (DEGs) across all cell types in ACG, and this remained the case when comparing control to PD or PDD clinical groups (FDR-adjusted p-value < 0.05, NACG = 1958, NIPL = 226, **Fig 2B**).

Similarly, we noted that of the two disease groups, PDD had the largest number of DEGs (FDR-adjusted p-value <0.05, Nup = 545, Ndown = 594) and this was consistent across both the ACG and IPL.

While these findings appeared to be consistent with a model of PDD as part of a continuum with PD, we found marked qualitative differences between the two clinical groups. More specifically, we identified differences in both the distribution of DEGs across the major cell types and the specific genes detected when comparing control to PD- and PDD-affected groups. The DEGs in PD were most frequently identified among excitatory neurons (FDR-adjusted p-value<0.05, CUX2_L2/3, Nup = 29, Ndown = 66; TLE4_L5/6, Nup = 48, Ndown = 69), inhibitory neurons (FDR-adjusted p-value < 0.05, RELN, Nup = 29, Ndown = 66) and astrocytes (FDR-adjusted p-value < 0.05, Astrocytes_CD44hi, Nup = 58, Ndown = 34). In contrast, the genes differentially expressed in PDD were more frequently found in glial populations, including two subtypes of astrocytes (FDR-adjusted p-value<0.05, Astrocytes_CD44hi, Nup = 103 Ndown = 84; Astrocytes SLC1A2hi_MERTKhi, Nup = 88, Ndown = 46) and microglia (FDR-adjusted p-value<0.05, Nup = 40, Ndown = 92) (**Fig 2C**). Furthermore, we observed a low overlap between DEGs detected in each disease group after accounting for cell type and direction of effect (**Fig 2D**). The cell types with the highest numbers of overlapping DEGs were oligodendrocytes derived from the IPL (25%), followed by pericytes (17%) and OPCs (16%) in the ACG.

Qualitative differences between PD and PDD were even more apparent when we conducted gene ontology and pathway enrichment analyses across the DEGs. We found clear differences in the numbers and types of gene ontology enrichments across the two clinical groups (**Fig 3E,F**). In fact, the only pathway which was enriched in the same cell type and with the same directionality in PD and PDD had the parent term calcium ion binding (sulfur compound binding, GO:1901681), which was upregulated in an astrocytic subtype (Astrocytes_SLC1A2hi_MERTKhi, 2.89×10^-2^ < FDR-adjusted p-value < 4.56×10^-2^). Focusing on PD alone, we identified pathway enrichments across almost all major cell types, with the highest numbers detected in astrocytes (FDR-adjusted p-value<0.05, N = 12 pathways), oligodendrocytes (FDR-adjusted padj<0.05, N= 8) and excitatory neurons (FDR-adjusted padj<0.05, N = 8). Notably, five pathways were identified in two or more cell types, namely molecular-adaptor activity, collagen containing extra-cellular matrix, heparin binding, the INO-80 type complex and the peri-ribosome subunit precursor. Astrocytes and oligodendrocytes were the two major cell types, which most commonly shared pathway enrichments (FDR-adjusted padj<0.05, N=4), suggesting a coordinated response to disease across macroglia. In contrast, pathway enrichments among DEGs in microglia tended to be unique to this cell type, and were relatively limited, including lipid biosynthetic process, regulation of immune system process, regulation of protein localisation and ribose phosphate metabolic process.

In PDD, pathway enrichments were primarily detected in only a few major cell types, with no enrichments detected in oligodendrocytes and OPCs, and surprisingly few changes in excitatory neurons given that this clinical group was distinguished by dementia. Microglia (FDR-adjusted p-value<0.05, N = 19 pathways) and astrocytes (FDR-adjusted p-value<0.05, N = 9 pathways) had the highest numbers of pathway enrichments. Interestingly, DEGs in both these major glial classes were enriched for terms associated with regulation of synaptic function, which is expected to be disrupted in dementia. Furthermore, when comparing pathway changes detected in PDD and PD, it was striking that some pathways were in fact inversely regulated. For example, while the lipid biosynthetic process was up-regulated in microglia in PD, it was down-regulated in astrocytes in PDD.

Finally, we assessed DEGs for evidence of enrichment for genes causally-implicated in disease, focusing on PD and AD given the high levels of co-pathology that exist particularly in PDD. We noted that all significant enrichments were detected in the ACG and in PDD. While DEGs detected in THEMIS_L5/6 neurons were enriched for genes causally-implicated in PD (FDR-adjusted p-value = 1.96 ×10^-2^), those in RORB_L4 neurons were enriched for genes causally-implicated in AD (FDR-adjusted p-value = 1.96 ×10^-2^) (**Fig 2G**). Thus, overall our analyses suggested that the molecular processes underlying PD and PDD differed not only in magnitude but also nature.

### Attenuation of regional identity is a feature of PDD, but not of PD

Cortical specialisation of neurons and glia across regions is well-established with some studies suggesting that with age and disease, such regional differences are attenuated (**Fig 3A**). To further explore this hypothesis in PD and PDD, we leveraged our study design and performed intra-individual regional analyses that compared features in the ACG and IPL in a pair-wise fashion. This statistically powerful approach provided us with regional differences between the ACG and IPL in the three clinical groups, namely controls, PD and PDD. In each case using the IPL as the baseline for comparison, we focused on three feature types, namely cell type proportions, gene expression and metabolic flux. As would be expected, in the control group we found several differences in cell type proportions in the ACG when compared to the IPL. Whereas RORB_L4 (FDR-corrected p-value = 3.37×10^-3^) and CUX2_L2/3 (FDR-corrected p-value = 4.03×10^-2^) excitatory neurons were less represented in the ACG than in the IPL, T-cells (FDR-corrected p-value = 6.91×10^-2^), venous endothelia (FDR-corrected p-value = 3.29×10^-2^) and smooth muscle cells (FDR-corrected p-value = 2.72×10^-2^) were all proportionally more represented in the ACG (**Fig 3B**).

In contrast in PD, many of these inherent regional differences were accentuated or sometimes even changed in directionality. For example, venous endothelia (FDR-corrected p-value > 5×10^-2^), smooth muscle cells (FDR-corrected p-value > 5×10^-2^) and T-cells (FDR-corrected p-value > 5×10^-2^) were in fact less represented in the ACG than IPL, potentially reflecting disease-associated changes in vascularisation. Furthermore, consistent with the significant decrease in RORB_L4 neurons in disease states in ACG, this specific class of neurons was particularly less represented in the ACG relative to IPL in PD (logFC = -5.71×10^-1^, FDR-corrected p-value = 2.45×10^-3^). This pattern was also apparent in PDD, where RORB_L4 (FDR-corrected p-value = 1.31×10^-2^) excitatory neurons dropped in proportion alongside CUX2_L2/3 excitatory neurons (FDR-corrected p-value = 1.87×10^-2^).

Next, we performed differential gene expression analysis comparing the ACG and IPL. In controls, excitatory neurons showed the most evidence for regional specialisation (**Fig 3C**) with RORB_L4 neurons having the highest number of DEGs (FDR-corrected p-value < 5×10^-2^, Ngenes = 7,232), followed by CUX2_L2/3 neurons (FDR-corrected p-value < 5×10^-2^, Ngenes = 5,970). In PD, this pattern of neuronal specialisation was largely maintained. However, in PDD there was a general reduction in differential gene expression, with neurons being particularly affected.

While RORB_L4 and CUX2_L2/3 neurons remained the most regionally distinct, we found that the numbers of DEGs were reduced to 3,077 and 1,947, respectively. Given that across clinical cohorts the highest numbers of differentially expressed genes were identified in neurons of the ACG samples, these findings indicate the loss of regional specialisation among excitatory neurons of this region rather than a shift in neuronal identity in the IPL. Notably, none of the PDD genes were enriched to specific pathways (**Fig 3E**).

Since neuronal activity is the major driver of metabolic state in the brain, the findings on neuronal specialisation in PD and PDD led us to investigate metabolic flux. Using single cell flux estimation analysis (scFEA), a method that has been validated in postmortem brain tissue^20^, we inferred changes in 22 metabolic supermodules, including fatty acid, aspartate metabolism and glycolysis and TCA cycle, across all cell types (**Fig 3D**). As expected, when comparing the ACG to IPL in the control group we found inherent differences in 15 metabolic supermodules with significant differences in up to five cell types. In PDD, there were fewer cell types with metabolic differences between regions than in controls. In contrast, in PD the differences observed in controls were amplified such that 21 supermodules had significant regional differences in at least one cell type. More specifically, we noted significant changes in flux within the glycolysis and TCA cycle across 10 cell types (FDR-corrected p-value <0.05), and in purine synthesis in nine cell types (FDR-corrected p-value <0.05). Furthermore, by focusing our analyses to specific cell types, we found that neurons were the most commonly affected by region-specific changes in metabolic flux (**Fig 3F**), and that glycolysis and the TCA cycle module was significantly disrupted in these cells (FDR-adjusted p-value < 0.05).

### RORB_L4 neurons have differential transcript usage in PDD

In the field of neurodegeneration there is increasing recognition of the importance of splicing and differential transcript use in driving disease processes. However this is challenging to study at a cell type-specific level. To address this issue, we studied 3’ untranslated region (UTR) use by cell type and across disease states, leveraging the relatively long read lengths generated for this data set, the high numbers of nuclei analysed per sample (5,850 nuclei) and a custom UTRome. More specifically, we used long-read RNA sequencing data from postmortem brain samples without neurodegenerative disease to generate a more accurate definition of 3’UTRs in the human brain, and adapted the scUTRquant pipeline for 3’ UTR quantification. This allowed us to generate single-cell 3’ UTR counts across all cell types, and infer changes in transcript use across clinical groups (**Fig 4A**). This analysis generated two major findings, namely that significant changes in transcript use by cell type and disease state occurred almost exclusively in excitatory neurons (88.9%), and that this signal distinguished between PDD and PD with 94.4% of all transcriptional changes detected in comparison of control and PDD clinical groups (**Fig 4B**).

**Figure 4.**
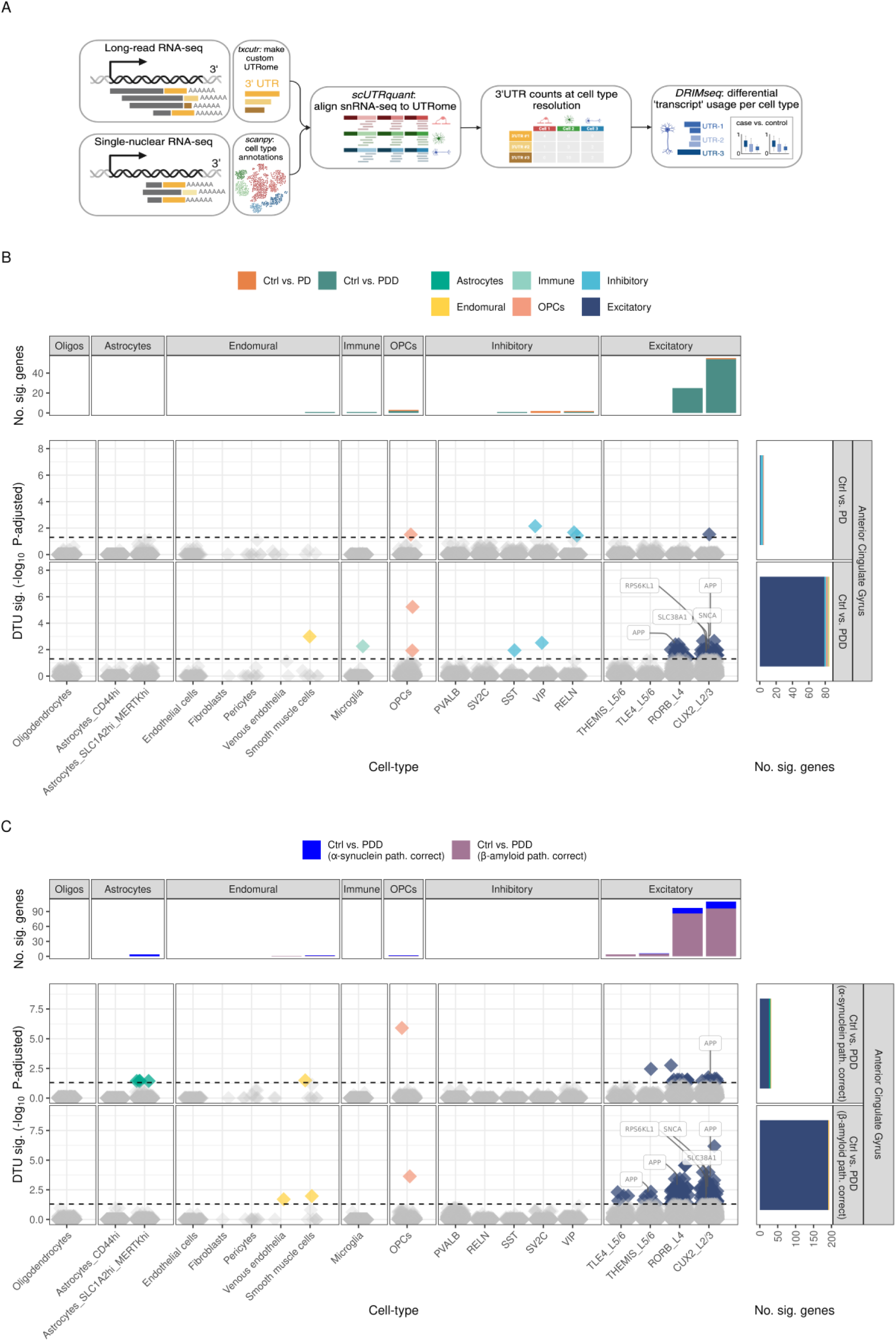
PDD has differential 3’ UTR usage which is enhanced when controlling for Aβ pathology. A. Workflow diagram of 3’ UTR usage analysis, which provides a proxy for transcript-level quantification at the level of cell types. This analysis required long-read RNA sequencing data for the generation of a custom reference 3’ UTR-ome. Single-nucleus RNA-sequencing data was aligned to the UTRome, providing a matrix of counts for each 3’ UTR per cell type. DRIMseq was finally used for evaluating differential 3’ UTR usage, followed by assessment of the transcripts associated with the differential 3’ UTRs. For the purpose of the protein domains contained within these transcripts of interest. B. Swarmplot of differential transcript usage significance levels (-log10 FDR-adjusted p-value) for each gene in each cell type. Genes highlighted in colour pass the significance threshold and gene name labels highlight PD-associated genes. This analysis was performed for control vs. PD, control vs PDD. C. Swarmplot of differential transcript usage significance levels (-log10 FDR-adjusted p-value) for each gene in each cell type. Genes highlighted in colour pass the significance threshold and gene name labels highlight PD-associated genes. Due to the presence of *APP* and *SNCA* among genes, we tested differential 3’ UTR use for control vs PDD with correction by Aβ and α-synuclein quantitative pathology to determine whether pathology impacted 3’ UTR use.

Across excitatory neuronal subtypes, we noted that the highest signals were observed in CUX2_L2/3 and RORB_L4 neurons, with 54 and 25 gene signals identified respectively. Furthermore, we found that among genes with evidence of differential transcript use in PDD, were genes causally implicated in PD and AD (N=7), most notably *SNCA* and *APP*. Given the high degree of variability in α-synuclein (Lewy bodies/mm2 = 2.19% ±SD2.66) and Aβ pathology (mean immunoreactivity = 2.42%±SD2.83) in our PDD cases, we wondered whether controlling for α-synuclein and Aβ load would reduce this signal and indicate a key role for this form of pathology in driving differential transcript usage. To test this hypothesis, we repeated the analysis, but included Aβ and α-synuclein pathological load in our modelling (**Fig 4C**).

Interestingly, we found that there were still significant changes in transcript use by cell type and disease state in both analyses. In the approach controlling for Aβ load, the overall number of significant signals increased by over 2-fold (N = 194), with 98.45% of the genes significant in excitatory neurons of which 10 genes were causally implicated in PD and AD. Furthermore, changes in 3’UTR use remained highly specific to excitatory neurons and PDD. When controlling for α-synuclein pathology there was a significant drop in the total number of transcriptomic changes (N = 32), but changes were still predominantly in excitatory neurons (81.25%). Differential transcript use of *APP,* but not *SNCA*, remained significant in this analysis when controlling for α-synuclein pathology.

Given that both *SNCA* and *APP* encode aggregating proteins of key importance in a range of neurodegenerative disorders, we studied differential transcript use in greater detail. Focusing on *SNCA*, we found that while there was no significant change in *SNCA* gene-level expression in CUX2_L2/3 excitatory neurons in PDD as compared to control groups, there was a significant change in the use of a specific 3’UTR. Interestingly, this 3’UTR is associated with a downstream open reading frame of 151 amino acids (ENST00000673902.1), rather than the major SNCA protein of 140 amino acids (ENST00000394991.8). In the case of *APP*, we focused on 3’UTR use in RORB_L4 neurons, again comparing PDD to control clinical groups. Although there was no evidence of differential expression of *APP* at the gene-level in this cell type (**Fig 5A**), we identified two 3’UTRs that were differentially used across clinical groups. While 3’ UTR #1 was used significantly more in controls than in PDD cases (FDR-adjusted p-value = 1.24×10^-03^), 3’ UTR #2 had the opposite direction of effect (FDR-adjusted p-value = 2.49×10^-05^) (**Fig 5C**). We noted that 3’ UTR #1 was associated with two transcripts (ENST00000439274 & ENST00000357903) encoding proteins of 714 and 751 amino acids respectively, while 3’ UTR #2 was associated with a single transcript (ENST00000448850) encoding a shorter open reading frame (485 amino acids) that did not include the Aβ 40/42 fragment (**Fig 5D**). Given that Aβ 40/42 fragments, generated through cleavage of APP, are the major constituent of Aβ plaques, we wondered whether the significant change in transcript use we observed could be a protective mechanism.

**Figure 5.**
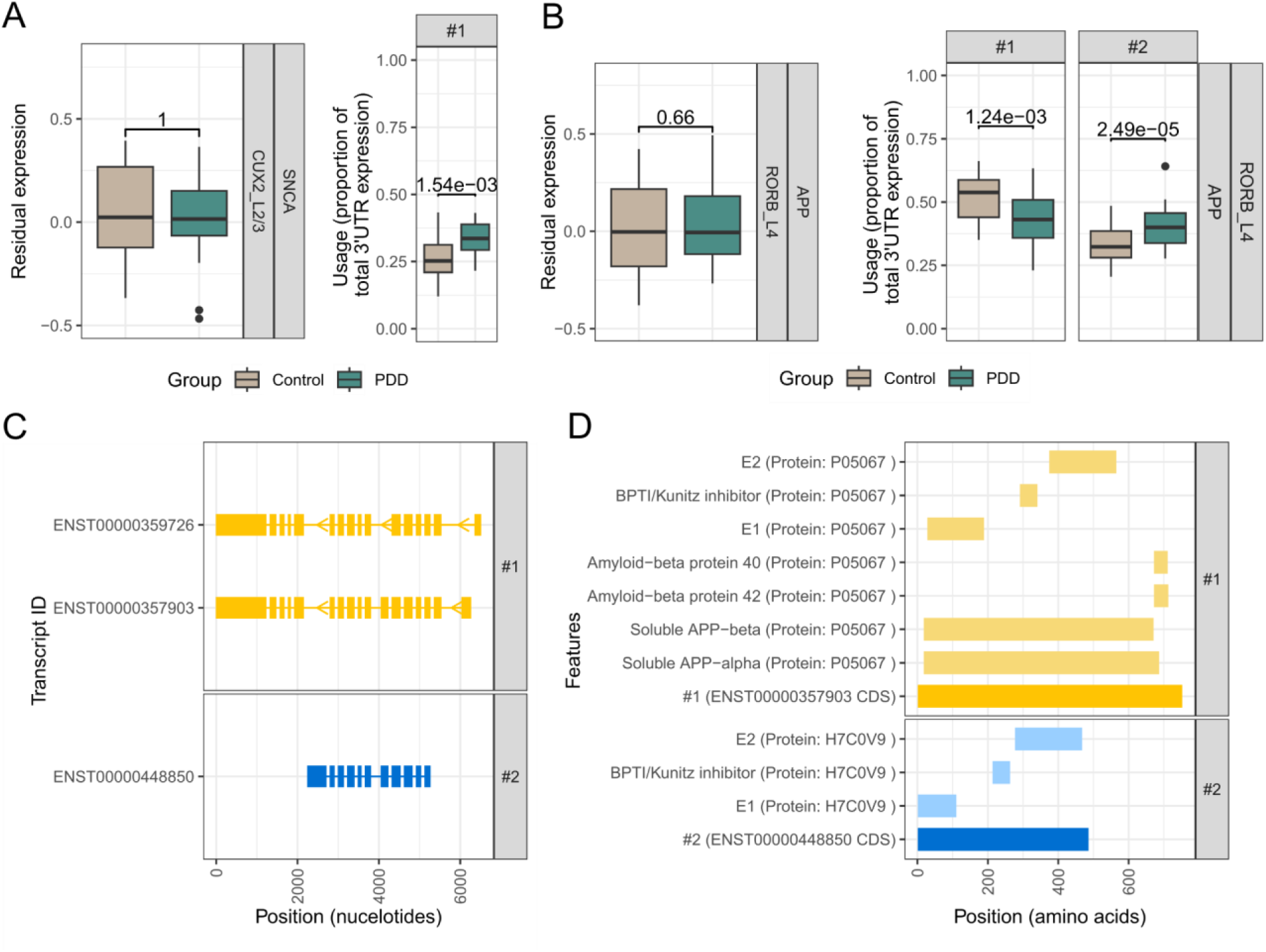
The two main 3’ UTRs of APP are differentially used in CUX2 and RORB excitatory neurons. A. Boxplots showing gene-level residual expression data and use of the top 3’ UTR for *SNCA* in CUX2_L2/3 excitatory neurons. Control and PDD distributions of *SNCA* gene-level residual expression were unchanged in control vs PDD, but there was a shift in 3’ UTR use when comparing both groups. 3’ UTR #1 was significantly more used in PDD when compared to controls (FDR-adjusted p-value = 1.54×10^-3^). B. Boxplots showing gene-level residual expression data and use of the top 3’ UTR for *APP* in RORB_L4 excitatory neurons. *APP* gene-level residual expression was unchanged when comparing control vs PDD, but there was a shift in 3’ UTR use when comparing both groups. 3’ UTR #1 was significantly less used in PDD when compared to controls (FDR-adjusted p-value = 1.24×10^-3^) and 3’ UTR #2 was significantly more used in PDD when compared to controls (FDR-adjusted p-value = 2.49×10^-5^). C. Transcript structure plots depicting the *APP* transcripts contained within each transcript bin, which is defined based on a unique 3’ UTR. We only show the transcripts for bins with significant differences when comparing control and PDD. D. Plots showing the protein coding sequence arising from transcripts in bins #1 and #2, in addition to protein chains and domains for the *APP* protein. One of the main differences in these 3’ UTRs was that 3’ UTR #2 (favoured in PDD) was associated with a shorter transcript, which does not include the pathology-associated Aβ protein 40/42 domain.

## Discussion

Despite its increasing prevalence, PD and PDD remain relatively understudied. Here, we addressed this knowledge gap and compared cellular composition, biological processes and molecular features in PD, PDD and neurotypical controls using the largest snRNAseq data for PDD generated to date. We found compelling evidence that PD and PDD are distinct at a cellular and molecular level with dysregulation of transcript use in excitatory neurons being a key distinguishing feature.

To date, only three studies have compared PD and PDD at a transcriptomic level in post-mortem human brain tissue^21–23^ and only one of these studies has done so using single cell profiling, making this analysis the largest of its kind. Our study identified differences between PD and PDD attributable to alterations in cell type proportions, cell type-specific gene expression and transcript use. While changes in the proportions of both neuronal and glial cell types were a feature of PD in the ACG, these were less evident in PDD and were generally subtle when compared to the compositional changes observed in AD^24^. Nonetheless, we noted that there was a significant fall in the proportion of RORB_L4 neurons in disease states and that, interestingly, this population of neurons also expressed high levels of genes causally-implicated in PD, including *SNCA*.

Both PD and PDD were characterised by extensive cell type-specific differential gene expression with evidence for the involvement of all major cell types at both disease stages. However, the specific genes and pathways impacted across these clinical groups showed a lower level of commonality than might have been expected. The majority of pathway enrichments were only identified on comparing single disease states to controls. Furthermore, we identified a small number of pathways which were enriched among genes up-regulated in PD but down-regulated in PDD. Given that this phenomenon was only observed in microglia, these findings raise the possibility of important phase-specific changes in inflammatory responses in PD progression that might trigger secondary pathologies. In fact, the enrichment of genes causally-implicated in AD among DEGs of RORB_L4 neurons in PDD provide some support for this hypothesis.

We explored the concept of disease phases further by leveraging data across the two analysed brain regions, namely the ACG and IPL. This form of within-individual analysis not only increased our power to detect disease-specific differences by effectively controlling for confounding variables, but also provided a broader understanding of the impact of disease on cortical specialisation. As expected, in neurotypical controls significant differences in cellular composition, gene expression and inferred metabolic activity were apparent. Although such regional differences appeared to be amplified in PD, this did not occur in PDD. In this clinical group there was an attenuation of regional specificity, which was most likely to be driven by effects on the ACG. While this requires further investigation, our analyses highlighted a potential role for the extracellular matrix in this process. We noted that whereas genes involved in collagen formation were upregulated in multiple cell types in PD, these pathways were down-regulated in PDD. In fact, the up-regulation of collagen in PD has been previously reported^25^, and is present in the context of traumatic brain injury or lesions in multiple sclerosis^26–28^.

Therefore, we think these findings may indicate processes that are occurring more globally in the PD brain and that warrant further investigation.

Of all cell types, neurons are the most regionally specialised, with increasing evidence to suggest they have a particularly complex transcriptome, characterised by high levels of alternative splicing. While 3’ snRNAseq does not enable the study of whole transcripts, it is an efficient means of studying 3’UTR use at a cell type-specific level. We made this approach more powerful in this study by leveraging long-read RNA-seq data from human brain samples to build an accurate brain-specific 3’UTRome. Remarkably, we found that significant changes in 3’ UTR use were largely detected in excitatory neurons (CUX2_L2L3 and RORB_L4), and that this was specific to PDD. Furthermore, amongst the genes showing the most significant differences in transcript use were *SNCA* and *APP*. Importantly, accounting for the Aβ pathological load, did not impact on these results and in fact, only increased gene detection. In contrast, when we controlled for Lewy body pathology, the majority of signals became non-significant. These findings strongly suggest that the transcriptomic changes we observed were driven primarily by Lewy body rather than Aβ pathology. This is important since not only is there growing interest and recognition of the importance of co-pathology across neurodegenerative disease, but with the development of new disease-modifying treatments there could also be implications for therapies in PDD. With this in mind, we conducted a detailed analysis of the relationship between 3’UTR use and open reading frames. Interestingly, in the case of *APP*, we noted that there was a reduction in the use of 3’UTRs associated with full-length *APP* transcripts in PDD. These transcripts encode proteins which can be cleaved to form Aβ 40- and 42-residuals, the major constituent of amyloid plaques^29^. Since we can only study surviving neurons this shift in transcript use could be a protective mechanism in PDD.

In summary, our study suggests that we should view Parkinson’s disease as a multi-phase disorder defined by distinct cellular and molecular phases of which one is dysregulation of RNA processing. Consequently, these results not only inform our understanding of the relationship between PD and PDD, but highlight the need to consider phase-specific therapies in PD.

## Acknowledgments

AFB, EKG, HM, JB, JE, JH, MGP, MR, NW, SG, ZJ were funded by Aligning Science Across Parkinson’s [Grant numbers: ASAP-000478 and ASAP-000509] through the Michael J. Fox Foundation for Parkinson’s Research (MJFF). AH was supported through the award of an Eisai-Leonard Wolfson Doctoral Training programme in Neurodegeneration. This work is supported by the UK Dementia Research Institute UKDRI-2206 through UK DRI Ltd, principally funded by the Medical Research Council. The authors wish to thank Carlo Sala Frigerio and Maria Rodriguez for support with protocols and troubleshooting snRNAseq generation, as well as Mervin M. Fansler’s support with running and adapting the scUTRquant pipeline. For the purpose of open access, the authors have applied a CC-BY public copyright license to all Author Accepted Manuscripts arising from this submission. RHR is currently employed by CoSyne Therapeutics (Lead Computational Biologist). All work performed for this publication was performed in her own time, and not as part of her duties as an employee. EKG has received paid consultancy from Isogenix limited within the last 12 months.

## Code availability

All sample preprocessing and analysis code will be made available upon publication: https://github.com/mgrantpeters/ASAP_teamhardy_snrnaseq

## Methods

### Brain sample selection

Samples were provided by the Queen Square Brain Bank. We examined samples from 64 donors, each of whom provided samples from two brain regions, namely the anterior cingulate cortex (ACG) and inferior parietal lobule (IPL). These brain regions were chosen to analyse regions affected by PD pathology at different stages of disease progression^30^. The 64 donors consisted of 27 females and 37 males and were categorised into 22 controls, 21 PDD cases and 21 PD cases (all at Braak stage 6). Full sample demographics are listed in Supplementary Table 1.

### Ethics

Informed consent to use the postmortem brain material for research is available for each donation. Human tissues were stored and analysed in accordance with UK legislation and the Human Tissue Authority license held by UCL Queen Square Institute of Neurology. The research project was approved by the NHS Health Research Authority, Ethics Committee London-Central.

### Tissue preparation

Brain tissues were flash-frozen at the time of post-mortem collection and stored at -80°C in the Queen Square Brain Bank. Upon retrieval, samples from the anterior cingulate gyrus and inferior parietal cortex were dissected on dry ice. To ensure anatomical consistency and pan-cortical representation across the cohort, all dissections were performed by a single neuropathologist (ZJ). Each sample comprised approximately 100 mg of full-thickness cortex with overlying leptomeninges.

### Neuropathology

Formalin-fixed and paraffin-embedded tissue blocks containing the cortex of the anterior superior and middle frontal gyri from the contralateral hemisphere were sectioned at 7μm on a manual rotary microtome (Leica RM2135) and immunostained for α-synuclein (MA1-90342; Thermo Scientific; 1:1500) and Aβ (M0872; Dako; 1:100) using Menarini or ROCHE Ventana Discovery automated staining platforms following manufacturer’s guidelines. The procedure employed biotinylated secondary antibodies, horseradish peroxidase-conjugated streptavidin complex and 3,3’-diaminobenzidine (DAB) as a chromogen and a haematoxylin counterstain. All immunostainings were performed with appropriate controls.

All slides were digitized on a Hamamatsu NanoZoomer S360 slide scanner with 40x objective (effective pixel size 0.23 µm/pixel). For figure preparation, representative images were taken on NZConnect (Hamamatsu), a web-based whole slide image (WSI) viewer. For quantitative image analysis, regions of interest (ROIs), namely the pancortex of the superior frontal and middle frontal gyri were manually annotated on NZConnect or NDP.view2 (Hamamatsu), an offline WSI viewer platform.

Annotations drawn on NZConnect were exported from the web-based database and downloaded as NDPA files using a custom Python script. Annotated WSIs were then imported into QuPath, v0.5.1^31^ for quantification by image analysis. Stain vectors for colour deconvolution were calculated from a composite image consisting of 1,000×1,000 pixels squares derived from 270 WSIs with 1% of extreme pixels ignored. This accounted for variability in haematoxylin and DAB staining intensities across slides and to establish stain vectors that are specific for the colour profile of the slide scanner. The mean deconvoluted DAB intensity within a representative ‘background’ annotation was measured and saved as a measurement value in their corresponding brain region annotation.

A threshold for positive DAB staining was calculated using the Otsu automatic threshold for hyperphosphorylated tau^32^, and the Triangle automatic threshold for Aβ ^33^. These calculated threshold values were added to the saved background DAB intensity measurement for each ROI. For Aβ, positive DAB staining area above the background-corrected threshold value was reported as a percentage area of immunoreactivity per region of interest.

WSIs stained for α-synuclein were analysed using QuPath with similar pre-processing steps to those described above, that included importing images and annotations and applying stain vectors for colour deconvolution. To detect Lewy bodies, the deep learning-based segmentation algorithm StarDist^34^ was used on the deconvoluted DAB channel. A random trees object classifier was trained from 1612 detection objects with 110 measurement features from a set of 40 training images to classify Lewy bodies from non-specific detections. The resulting Lewy body object classifier was then applied to the entire cohort of WSIs and results reported as Lewy bodies/mm^2^.

### Single-nucleus RNA preparation and sequencing

Nuclei were isolated from frozen brain tissue in preparation for single-nucleus RNA-sequencing using a protocol described here: https://dx.doi.org/10.17504/protocols.io.yxmvm25xng3p/v1. Briefly, ∼100mg frozen tissue per sample was homogenized using a dounce tissue grinder with a custom homogenization buffer. Samples were then layered onto a density gradient medium and centrifuged in an Optima XPN100 ultracentrifuge at 7700 rotations per minute (RPM) for 30 minutes at 4◦C in an SW 41 Ti swinging-bucket rotor (Beckman Coulter). The gradient was removed and nuclei were resuspended and filtered twice to remove any further debris. The samples were then counted on a LUNA-FL dual fluorescence cell counter (Logos Biosystems) and diluted to the concentration required to capture a target of 8000 nuclei per sample. Samples were processed using 10X Genomics GEM isolation technology with the Chromium accessory, and complementary DNA (cDNA) amplification and library construction was performed using the Single Cell 3’ Reagent Kits, as per manufacturer’s instructions. Amplified cDNA and subsequent cDNA libraries were quantified on a QUBit 4 fluorometer (ThermoFisher) and the distribution of molecule sizes determined using the Agilent 4200 Tapestation (Agilent). Pooled libraries were loaded onto S4 flow cells and sequenced using the Novaseq 6000 Sequencing System (Illumina).

### snRNAseq data preparation with panpipes

We used a modified version of the dev branch of the nf-core scrnaseq nextflow pipeline (PMID: 32055031, https://zenodo.org/records/6656322). More specifically, we modified v2.0.1 dev (commit: #0104519), and added an option to make matrix conversion optional (modification detailed [here](https://github.com/RHReynolds/nf-core-scrnaseq/pull/1)).

Reads were mapped to the GRCh38 human reference genome via STARsolo (v 2.7.8a) using gene annotations from Ensembl v107 (PMID: 23104886, 29155950; bioRxiv: 2021.05.05.442755). Per-sample 2-pass mapping was used, wherein two rounds of mapping were performed to improve the sensitivity of novel splice junction detection. ENCODE standard options for long RNA-seq were used (as detailed in STAR manual, v 2.7.8a), with the exception of (i) --outFilterMultimapNmax, which was set to 1, thus retaining only uniquely mapped reads and (ii) --alignSJDBoverhangMin, which was set to the STAR default of a minimum 3 bp overhang required for an annotated spliced alignment. In addition, to generate a filtered gene/cell count matrix almost identical to CellRanger’s, parameters were set as described in the STARsolo documentation (commit: [#fb84afe](https://github.com/alexdobin/STAR/commit/fb84afe46ef590eb90e32297106bf62818a0495f)).

Aligned data were ingested to panpipes for QC, preprocessing and clustering^35^. Nuclei were filtered based on whether they passed conventional QC metrics, namely, doublet detection score with scrublet (score<0.15), percentage of mitochondrial transcripts in RNA counts (<5%), percentage of ribosomal transcripts in RNA counts (<5%). This resulted in a total of 789,708 high quality nuclei. Batch correction was performed across samples using Harmony^36^ and community detection with the leiden algorithm. Major cell types were detectable with cell type markers (Astrocytes: AQP4, GFAP; Endomural cells: CLDN5; Neurons: GABRB2; Inhibitory neurons: GAD2; Immune cells: CD74, PTPRC, TREM2, APOE; Oligodendrocytes: PLP1, MBP; OPCs: PDGFRA, BCAS1). These major cell types were then subclustered further, similarly to other single nucleus RNA sequencing approaches^24,37,38^. For this, each cell type was batch corrected separately using Harmony again and community detection was performed again and leiden community detection was used to partition UMAP dimensionality-reduced clusters. Using clustree cluster hierarchy, leiden resolutions were chosen per cell type based on clustering stability.

Cluster annotations were performed using a combination of cell markers and the highly variable genes calculated for each cluster, as implemented by scanpy^39^. Based on a combination of highly variable genes and marker genes, cell types were annotated across 4 levels of increasing resolution, from level 0 (major cell type) to level 3 (high resolution cell type/state).

snRNA-seq data were pseudobulked for downstream analyses. The aggregateToPseudoBulk function from the dreamlet R package (version 0.99.16)^19^ was used to sum gene expression counts of nuclei by participant ID, brain region (IPL, ACG) and cell type annotation. This was performed separately for each annotation level.

### Selection of covariates

Covariates driving significant variation in the gene expression data were discovered using an unbiased, data-driven approach. First, all technical (sample preparation, nuclei extraction, sequencing, mapping), clinical, and histopathological metadata were combined. Uninformative or unusable variables were removed, including IDs, those with zero variance and those that were a proxy for disease group. We then performed a second tier of filtration, removing variables with high missingness and selecting one from variable pairs with high collinearity. Of the variables remaining, those with missing values underwent mean imputation (for numericals), or common value imputation (for categoricals). For assessing pairs of continuous variables, collinearity was assessed using the caret R package (version 6.0-94), with a Spearman’s rank correlation coefficient cut-off of set at 0.7. For every correlation above this threshold, the variable with the lowest average correlation was kept and the other covariate removed. To assess collinearity of pairs of categorical variables, a χ-squared test was used. For pairs of numeric and categorical variables, we used the Kruskal-Wallace test and for pairs of numeric variables we used a Spearman’s rank correlation test. The resulting P-values were assessed using a significance threshold adjusted for multiple comparisons (P<0.05/(number of tests)).

Variables were then removed from pairs that demonstrated a significant categorical/categorical or categorical/continuous relationships. The contribution to variance in gene expression of the remaining variables was assessed by two methods: (1) variancePartition^40^; variancePartition R package) and (2) Principal Component Analysis (PCA; prcomp R package). VariancePartition assesses the contribution of variance of putative covariates at the gene level, whereas PCA-based dimensionality reduction generates per-sample eigenvectors which can then be correlated with putative covariates to assess their relative importance. We employed these two methods to determine our final covariates, recognising the need to control not only for covariates with global impacts on expression, but also those that may confound the expression of small subsets of genes. Putative covariates were then assessed against three metrics: (1) variancePartition: the maximum variance explained percentage, providing an assessment of whether a covariate comprises a large proportion of the variation of any gene(s), (2) variancePartition: the 3rd quartile of the variance explained distribution, providing an assessment of whether a covariate comprises a sizeable proportion of the variation of a larger proportion of genes and (3) the Spearman’s rank correlation coefficient between PC-X and covariate. For each metric, data-driven cut-offs were set using visualisation and cross-cell type concordance was assessed. Covariates passing the cut-offs in >50% (4/7) of cell types were selected as the final covariates. This resulted in the selection of the following five covariates: sex, RIN, deletion length, insertion length and uniquely mapped percent.

### Cell type proportions

For the cross-group analysis, differences in cell type proportions across groups were assessed using crumblr (https://github.com/GabrielHoffman/crumblr/). Briefly, occurrences of each cell type in each sample were summed. The data was covariate corrected (variables: deletion_length, insertion_length, rin_bxp, sex and uniquely_mapped_percent) using dream and the fit was smoothed using eBayes(). This comparison was performed separately for each brain region (ACG, IPL) and pairwise comparisons were performed for controls vs PD and controls vs PDD. Finally, p-values were FDR-corrected for each cell type compartment.

### Outlier detection

Following pseudobulking, we performed outlier detection. The outlier detection method employed was adapted from^41^ and utilised pseudobulked, normalised (log2 Fragments Per Million (FPM)), covariate corrected expression values. In brief, a sample was considered an outlier if it (1) had an absolute Z-score greater than 3 for any of the top 10 principal components and (2) had a sample connectivity score of less than -2 and (3) fulfilled the first two criteria in at least 50% (4/7) of cell types. Sample connectivity was calculated using the WGCNA R package^42^. Using this method, we identified one outlier.

### Differential gene expression analysis

Differential expression analysis was carried out using the Dreamlet R package^19^ version 0.99.16^19^. First, pseudobulked expression values were normalised (per cell type) using the dreamlet::processAssays function, setting the ‘min.count’ argument (minimum number of reads for a gene to be considered expressed in a sample) to 10. The resulting normalised counts were then passed to the dreamlet::dreamlet function to perform differential expression. Six contrasts were defined, setting the comparisons that would be made (all possible comparisons between three sample groups (PD, PDD, Control) for two tissues (ACG, IPL). Also supplied to the dreamlet function were normalised expression values and the following formula that included five covariates:

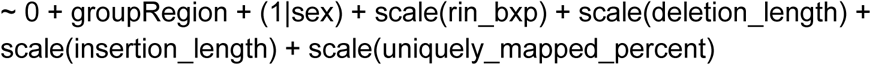

Between-tissue differential expression was performed similarly, though using different contrasts and a slightly altered design formula. For this analysis, three contrasts were defined, setting the comparisons between two tissues (ACG, IPL) within each of the three disease groups (PD, PDD, Control). In the between-group analysis, (1|sex) accounted for inter-individual variability due to sex-related differences. However, for the between-tissue analysis, a paired design was used, where each individual contributed samples from both ACG and IPL. In this case, (1|patient) was incorporated as a random effect to explicitly model inter-individual variability, ensuring that differences between ACG and IPL were assessed within each individual rather than across individuals. As such, the following formula was supplied to the dreamlet function:

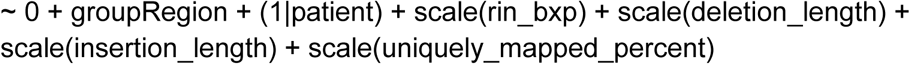

### Pathway enrichment

Pathway enrichment analysis was performed using the clusterProfiler R package (version 4.8.2) to identify biological pathways significantly enriched in the differentially expressed genes. Gene Ontology (GO), Kyoto Encyclopedia of Genes and Genomes (KEGG) and Reactome pathway databases were used as references for enrichment. Prior to analysis, gene IDs were converted to Entrez IDs using the AnnotationDbi package (version 1.62.2). The enrichGO and enrichKEGG functions from clusterProfiler as well as the enrichPathway function from ReactomePA (version 1.44.0) were applied to identify overrepresented terms and pathways in the set of differentially expressed genes. The model passed to the geneClusters argument was as follows:

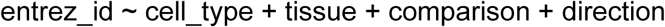

The analysis was run for each tissue (i.e. ACG), cell type (i.e. excitatory neurons) and comparison (i.e. PD vs. Control) separately. For each run, the set of all genes expressed in a given tissue and cell type was used as the background gene set for enrichment, supplied to the ‘universe’ argument. The results were adjusted for multiple testing, with terms and pathways showing an adjusted p-value of < 0.05 considered statistically significant. Adjustment of P-values was performed separately for each cell_type/tissue/comparison run of the analysis.

### Familial, GWAS and PGRS implicated PD gene enrichment

To enable disease gene list enrichment analyses, we curated gene panels using publicly available datasets and standardised filtering criteria. Our approach incorporated genes implicated in Parkinson’s disease (PD) and Alzheimer’s disease (AD), considering both rare Mendelian disease-associated genes and common variant GWAS-implicated genes. For the PD rare Mendelian gene list, we utilised two independent sources. The first was Blauwendraat et al. (2020), from which we extracted genes listed in Table 1 of the manuscript, retaining only those classified as high or very high confidence for PD causation. The second source was OMIM, where we performed a keyword search for “Parkinson disease” and retained only genes with strong disease associations. Genes with weak, disproven, or uncertain evidence (GIGYF2, UCHL1, EIF4G1) were manually excluded. For AD, rare Mendelian genes were sourced from OpenTargets, filtering for genes associated with Alzheimer’s disease and requiring a genetic association score >0.8.

For the common variant GWAS-implicated gene lists, we utilised the most recent and comprehensive datasets for each disease. For PD, we incorporated genes identified in the multi-ancestry GWAS by Kim et al. (2023). For AD, we used the Bellenguez et al. (2022) GWAS dataset. To the latter list, APOE was manually added, as it was excluded from the original study. Gene symbols were mapped to Ensembl gene IDs using the org.Hs.eg.db R package, with manual curation applied to resolve missing or ambiguous identifiers.

### ACG/IPL paired cell type proportion analysis

We calculated the cell type proportions for all cell types from each sample. The proportions in ACG and IPL were compared using a paired method, meaning that any individuals that did not have samples representing both regions were excluded from this analysis. Significance was calculated using a paired Wilcoxon test, as implemented in the python library scipy. The p-values were FDR-corrected using the fdrcorrection function from the python library statsmodels. The median cell type proportions per cell type per group were calculated and the ratio of the ACG median relative to the IPL median were taken, yielding the logFC in ACG for ease of visualisation. These results were visualised using the python seaborn library clustermap function.

### ACG/IPL paired metabolic inference from transcriptomic data

Metabolic flux was inferred from transcriptomic data using single cell flux estimation analysis (scFEA)^20^. Pseudobulked gene expression data was CPM normalised and log transformed, then ingested to scFEA. Matching the method used by the Seurat function RegressOut, the metabolite data was fit to a linear regression model using R (lm()). Statistical analysis comparing disease groups was performed in a pair-wise fashion with the Wilcoxon test and multiple testing corrected using FDR.

### Generation of custom UTRome

To construct a custom UTRome for downstream analyses, the GTF file from the long-read RNA was processed as follows. The GTF file was filtered to retain transcripts on standard chromosomes (chr1-22, X, Y), exclude ambiguous PAR regions, and remove transcripts with undefined or ambiguous strand information. This filtering step was performed using rtracklayer (version 1.60.1) and plyranges (version 1.20.0) in R. The cleaned GTF file was processed using SQANTI3 (version 5.3.0) to identify open reading frames (ORFs) and classify transcripts.

Transcripts were filtered using the default SQANTI3 rules to remove low-quality or potentially artifactual isoforms. Transcripts were further filtered to include only those associated with protein-coding genes, as defined by GENCODE. This was achieved by cross-referencing SQANTI3 classifications with GENCODE’s “protein_coding” gene and transcript annotations. Only transcripts classified as “coding” by SQANTI3 and annotated as “protein_coding” in GENCODE were retained. The filtered transcriptome was prepared for scUTRquant input using txcutr (version 1.6.0). A TxDb object was created from the filtered GTF file using GenomicFeatures (version 1.52.2), and truncation was applied with the truncateTxome function.

### Transcript-level quantification and differential transcript usage analysis

Single-nucleus transcript-level quantification was performed using scUTRquant^43^. Reads were pseudoaligned to a 3’ UTR reference, quantifying unique 3’ UTRs for each gene. We then utilised this quantification as a proxy for transcript-level quantification. Fastq files, sample ID mapping, cell type annotations (as described previously) and a custom UTRome were supplied to the scUTRquant Snakemake pipeline. The pipeline (per-sample) outputs were then aggregated and the nuclei were filtered to remove cells with very low transcript diversity (effective transcripts >=300). The effective transcript metric is based on Shannon entropy and was calculated utilising the compute_effective_count from the scUTRquant GitHub repository. Normalised counts were then calculated using the scuttle package (version 1.10.3)^44^.

Pseudobulking was then performed using the dreamlet aggregateToPseudoBulk function as previously described. To derive differential transcript usage, we implemented a Dirichlet Multinomial model using the DRIMSeq R package (version 1.28.0)^45^. To ensure minimum reliability of genes and transcripts analysed, DRIMSeq implements a filtration step using the dmFilter function. The values supplied to each argument were as follows.

Min_samps_gene_expr was set to 75% of the size of the smallest group, min_samps_feature_expr was set to the size of the smallest group and both min_gene_expr and min_feature_expr were set to 10. The same covariates were used as used for the differential gene expression analysis and as such the following model was supplied to DRIMSeq:

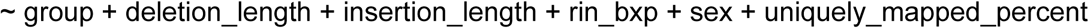

Staging was then used to provide appropriate FDR control using the stageR package (version 1.22.0)^46^. DRIMSeq was run in this way for each tissue (ACG, IPL), for each cell type and for each comparison (PDD vs. Control, PD vs. Control). Differential transcript usage was visualised using ggplot2 (version 3.5.1) and custom scripts.

For differential transcript usage controlling for neuropathology measures, the same protocol as above was used, but in these instances, the model was adjusted to include continuous measures of the relevant pathology. To control for alpha-synuclein pathology, Lewy Bodies per mm^2^ in the frontal cortex of the relevant individual was used and to control for amyloid-beta pathology, the % area of amyloid-beta immunoreactivity in the frontal cortex of the relevant individual was used.

### APP and SNCA transcript characterisation and visualisation

Differential transcript usage (DTU) for key genes (*APP* and *SNCA*) was characterised and visualised using a two-step analysis. DRIMseq gene-level –log₁₀ P-values were visualised as dot plots for each comparison, cell types and tissue. Distinct peaks in these plots, corresponding to genes with highly significant DTU, prompted further investigation of *APP* and *SNCA*. For these genes, case–control expression distributions were plotted at both the transcript bin and gene levels to ascertain whether transcript-level changes occurred independently of overall gene expression changes. To further understand the biological relevance of the buns, it was important to determine which individual transcripts contributed to each 3′UTR bin. To this end, the scUTRquant output merge file (*.merge.tsv) was interrogated, allowing the mapping of transcript identifiers to 3′UTR bins. To investigate potential protein-level consequences of the DTU events, protein domain information was retrieved for the transcripts representing each bin via Ensembl and UniProt REST APIs.

### Oxford Nanopore Sequencing

Six post-mortem brain samples were collected from the frontal cortex of control brains. Approximately 50-100 g of frozen tissue was used per brain sample for RNA extraction. Tissue lysis was performed using QIAzol and RNA was extracted using the RNeasy 96 Kit (Qiagen) with an on-membrane DNase treatment, as per manufacturer instructions. Samples were quantified by absorption on the QIAxpert (Qiagen), and RNA integrity number (RIN) measured using the Agilent 4200 Tapestation (Agilent). Libraries were prepared for Oxford Nanopore PromethION Sequencing using the standard SQK-PCS111 cDNA-PCR Sequencing Kit. 200ng total RNA was used as input per sample and 12 cycles were performed during the amplification step. Approximately 25 mol of each sample was loaded onto a separate flow cell before sequencing on the Oxford Nanopore PromethION. Flow cells were refuelled at 65 hours and runs were terminated at approximately 92 hours.

### GTF File Generation

Libraries were basecalled as standard with the Oxford Nanopore High-accuracy model. NanoStat (1.6.0) and Pychopper (2.7.6) were run on each library. Reads were then mapped using Minimap2 (2.26). Stringtie (2.2.1) was used to make an initial .gtf file of each library using Gencode annotation (v.38). These files were then merged using Stringtie to make a total gtf, and Gffcompare (0.12.6) was used to compare the bulk file with the Gencode v38 annotation to generate class codes.

## Bibliography

1. Mayeux, R. et al. A population-based investigation of Parkinson’s disease with and without dementia. Relationship to age and gender. Arch. Neurol. 49, 492–497 (1992).

2. Fink, A., Dodel, R., Georges, D. & Doblhammer, G. The Impact of Sex-Specific Survival on the Incidence of Dementia in Parkinson’s Disease. Mov. Disord. 38, 2041–2052 (2023).

3. Reid, W. G. et al. A longitudinal of Parkinson’s disease: clinical and neuropsychological correlates of dementia. J. Clin. Neurosci. 3, 327–333 (1996).

4. Hely, M. A., Reid, W. G. J., Adena, M. A., Halliday, G. M. & Morris, J. G. L. The Sydney multicenter study of Parkinson’s disease: the inevitability of dementia at 20 years. Mov. Disord. 23, 837–844 (2008).

5. McKeith, I. G. et al. Diagnosis and management of dementia with Lewy bodies: Fourth consensus report of the DLB Consortium. Neurology 89, 88–100 (2017).

6. Emre, M. et al. Clinical diagnostic criteria for dementia associated with Parkinson’s disease. Mov. Disord. 22, 1689–707; quiz 1837 (2007).

7. Robinson, J. L. et al. Pathological combinations in neurodegenerative disease are heterogeneous and disease-associated. Brain 146, 2557–2569 (2023).

8. Schneider, J. A., Arvanitakis, Z., Bang, W. & Bennett, D. A. Mixed brain pathologies account for most dementia cases in community-dwelling older persons. Neurology 69, 2197–2204 (2007).

9. Josephs, K. A. et al. TDP-43 is a key player in the clinical features associated with Alzheimer’s disease. Acta Neuropathol. 127, 811–824 (2014).

10. Brockmann, K. et al. GBA-associated Parkinson’s disease: reduced survival and more rapid progression in a prospective longitudinal study. Mov. Disord. 30, 407–411 (2015).

11. Papadimitriou, D. et al. Motor and Nonmotor Features of Carriers of the p.A53T Alpha-Synuclein Mutation: A Longitudinal Study. Mov. Disord. 31, 1226–1230 (2016).

12. Wu, L. Y. et al. Investigation of the genetic aetiology of Lewy body diseases with and without dementia. Brain Commun. 6, fcae190 (2024).

13. Real, R. et al. Association between the LRP1B and APOE loci and the development of Parkinson’s disease dementia. Brain 146, 1873–1887 (2023).

14. Chia, R. et al. Genome sequencing analysis identifies new loci associated with Lewy body dementia and provides insights into its genetic architecture. Nat. Genet. 53, 294–303 (2021).

15. Lennon, V. A. et al. A serum autoantibody marker of neuromyelitis optica: distinction from multiple sclerosis. Lancet 364, 2106–2112 (2004).

16. O’Connor, K. C. et al. Self-antigen tetramers discriminate between myelin autoantibodies to native or denatured protein. Nat. Med. 13, 211–217 (2007).

17. Solomon, A. J. et al. Differential diagnosis of suspected multiple sclerosis: an updated consensus approach. Lancet Neurol. 22, 750–768 (2023).

18. Blauwendraat, C., Nalls, M. A. & Singleton, A. B. The genetic architecture of Parkinson’s disease. Lancet Neurol. 19, 170–178 (2020).

19. Hoffman, G. E., et al. Efficient differential expression analysis of large-scale single cell transcriptomics data using dreamlet. BioRxiv (2024) doi:10.1101/2023.03.17.533005.

20. Alghamdi, N. et al. A graph neural network model to estimate cell-wise metabolic flux using single-cell RNA-seq data. Genome Res. 31, 1867–1884 (2021).

21. Irmady, K. et al. Transcriptome patterns and their regulation by alternative splicing in Parkinson’s disease. (P12-11.006). Neurology 98, (2022).

22. Feleke, R. et al. Cross-platform transcriptional profiling identifies common and distinct molecular pathologies in Lewy body diseases. Acta Neuropathol. 142, 449–474 (2021).

23. Henderson-Smith, A. et al. Next-generation profiling to identify the molecular etiology of Parkinson dementia. Neurol. Genet. 2, e75 (2016).

24. Mathys, H. et al. Single-cell atlas reveals correlates of high cognitive function, dementia, and resilience to Alzheimer’s disease pathology. Cell 186, 4365–4385.e27 (2023).

25. Raghunathan, R., Hogan, J. D., Labadorf, A., Myers, R. H. & Zaia, J. A glycomics and proteomics study of aging and Parkinson’s disease in human brain. Sci. Rep. 10, 12804 (2020).

26. Bradbury, E. J. & Burnside, E. R. Moving beyond the glial scar for spinal cord repair. Nat. Commun. 10, 3879 (2019).

27. Hu, X. et al. Spinal cord injury: molecular mechanisms and therapeutic interventions. Signal Transduct. Target. Ther. 8, 245 (2023).

28. Mohan, H. et al. Extracellular matrix in multiple sclerosis lesions: Fibrillar collagens, biglycan and decorin are upregulated and associated with infiltrating immune cells. Brain Pathol. 20, 966–975 (2010).

29. Gu, L. & Guo, Z. Alzheimer’s Aβ42 and Aβ40 peptides form interlaced amyloid fibrils. J. Neurochem. 126, 305–311 (2013).

30. Rietdijk, C. D., Perez-Pardo, P., Garssen, J., van Wezel, R. J. A. & Kraneveld, A. D. Exploring braak’s hypothesis of parkinson’s disease. Front. Neurol. 8, 37 (2017).

31. Bankhead, P. et al. QuPath: Open source software for digital pathology image analysis. Sci. Rep. 7, 16878 (2017).

32. Otsu, N. A Threshold Selection Method from Gray-Level Histograms. IEEE Trans. Syst. Man Cybern. 9, 62–66 (1979).

33. Zack, G. W., Rogers, W. E. & Latt, S. A. Automatic measurement of sister chromatid exchange frequency. J. Histochem. Cytochem. 25, 741–753 (1977).

34. Schmidt, U., Weigert, M., Broaddus, C. & Myers, G. Cell Detection with Star-Convex Polygons. in Medical Image Computing and Computer Assisted Intervention – MICCAI 2018: 21st International Conference, Granada, Spain, September 16-20, 2018, Proceedings, Part II (eds. Frangi, A. F., Schnabel, J. A., Davatzikos, C., Alberola-López, C. & Fichtinger, G.) vol. 11071 265–273 (Springer International Publishing, 2018).

35. Curion, F. et al. Panpipes: a pipeline for multiomic single-cell and spatial transcriptomic data analysis. Genome Biol. 25, 181 (2024).

36. Korsunsky, I. et al. Fast, sensitive and accurate integration of single-cell data with Harmony. Nat. Methods 16, 1289–1296 (2019).

37. Jäkel, S. et al. Altered human oligodendrocyte heterogeneity in multiple sclerosis. Nature 566, 543–547 (2019).

38. COvid-19 Multi-omics Blood ATlas (COMBAT) Consortium. A blood atlas of COVID-19 defines hallmarks of disease severity and specificity. Cell 185, 916–938.e58 (2022).

39. Wolf, F. A., Angerer, P. & Theis, F. J. SCANPY: large-scale single-cell gene expression data analysis. Genome Biol. 19, 15 (2018).

40. Hoffman, G. E. & Schadt, E. E. variancePartition: interpreting drivers of variation in complex gene expression studies. BMC Bioinformatics 17, 483 (2016).

41. Gandal, M. J. et al. Broad transcriptomic dysregulation occurs across the cerebral cortex in ASD. Nature 611, 532–539 (2022).

42. Zhang, B. & Horvath, S. A general framework for weighted gene co-expression network analysis. Stat. Appl. Genet. Mol. Biol. 4, Article17 (2005).

43. Fansler, M. M., Mitschka, S. & Mayr, C. Quantifying 3’UTR length from scRNA-seq data reveals changes independent of gene expression. Nat. Commun. 15, 4050 (2024).

44. McCarthy, D. J., Campbell, K. R., Lun, A. T. L. & Wills, Q. F. Scater: pre-processing, quality control, normalization and visualization of single-cell RNA-seq data in R. Bioinformatics 33, 1179–1186 (2017).

45. Nowicka, M. & Robinson, M. D. DRIMSeq: a Dirichlet-multinomial framework for multivariate count outcomes in genomics. [version 2; peer review: 2 approved]. F1000Res. 5, 1356 (2016).

46. Van den Berge, K., Soneson, C., Robinson, M. D. & Clement, L. stageR: a general stage-wise method for controlling the gene-level false discovery rate in differential expression and differential transcript usage. Genome Biol. 18, 151 (2017).

